## Supplementary figures and images for "The Rab7-Epg5 and Rab39-ema modules cooperatively position autophagosomes for efficient lysosomal fusions"

### Supplemental Data 1

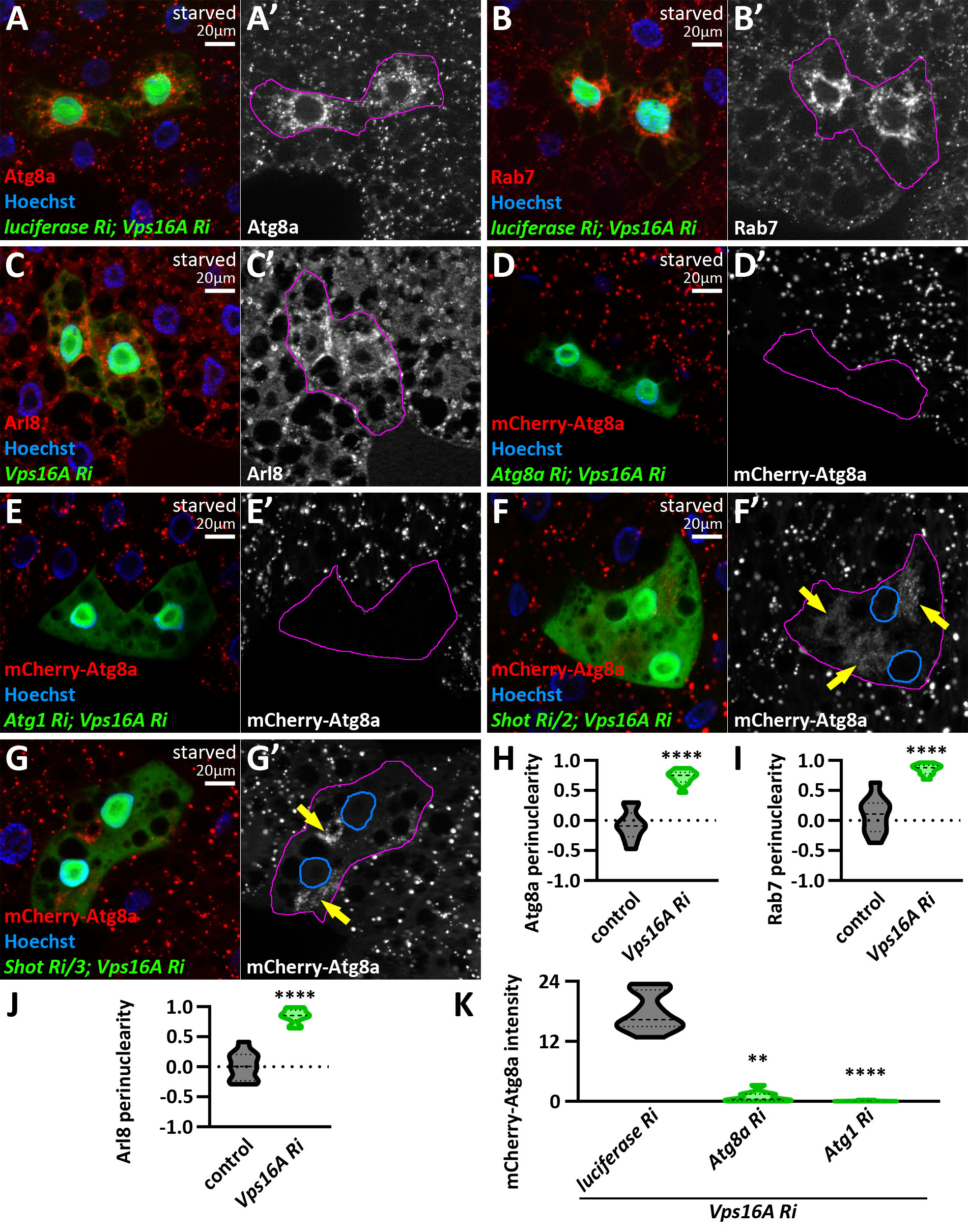

### Supplemental Data 2

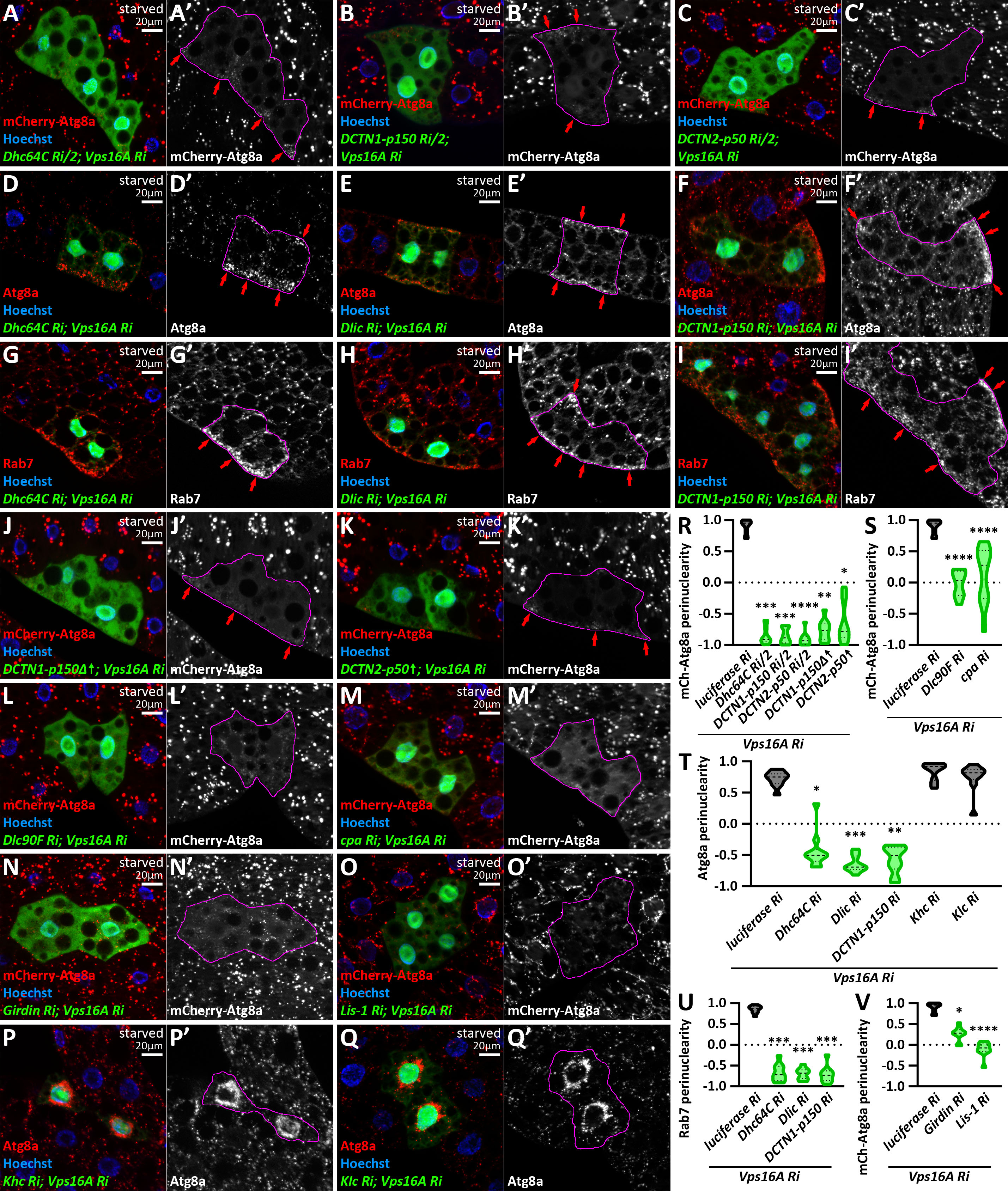

### Supplemental Data 3

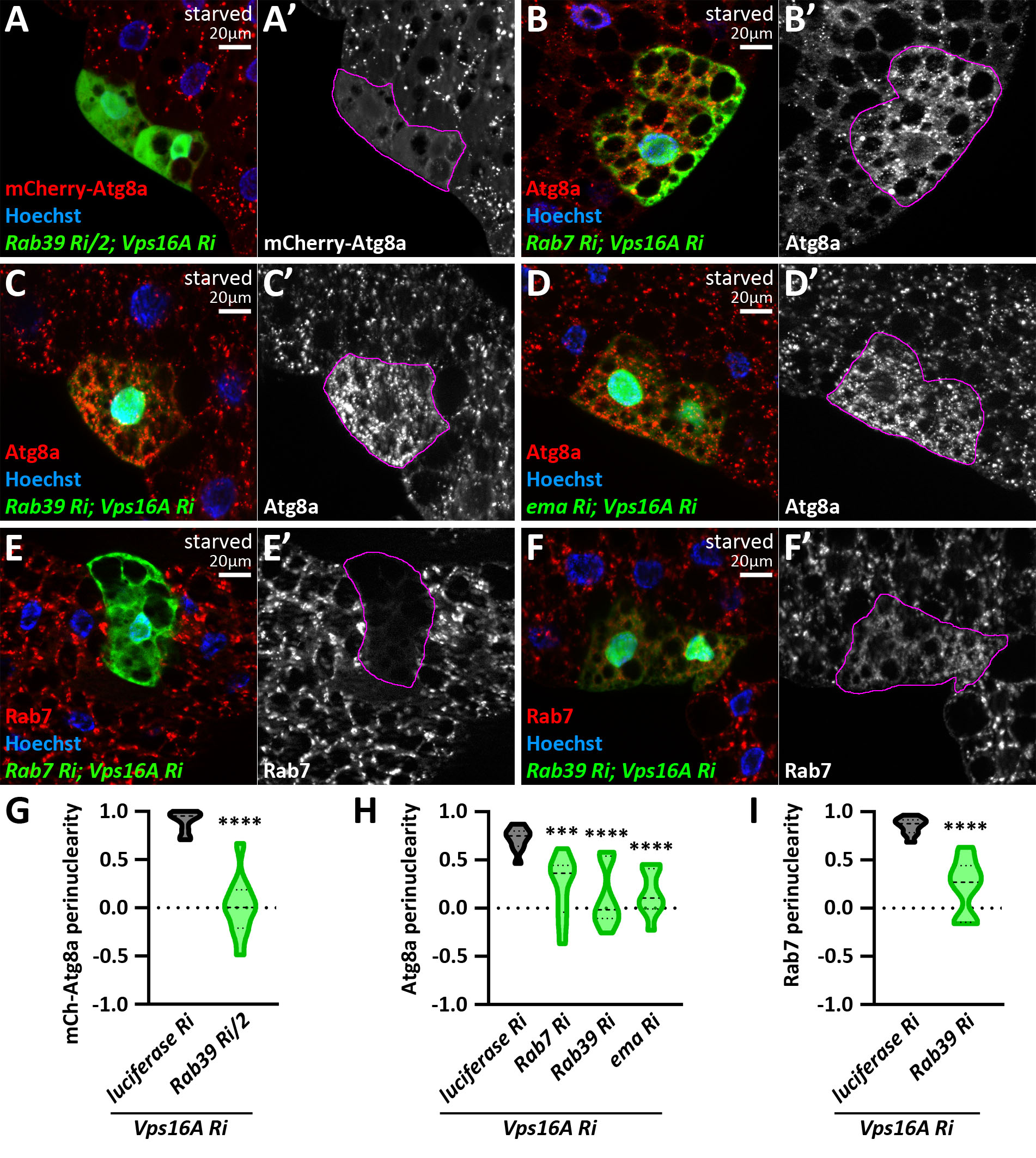

### Supplemental Data 4

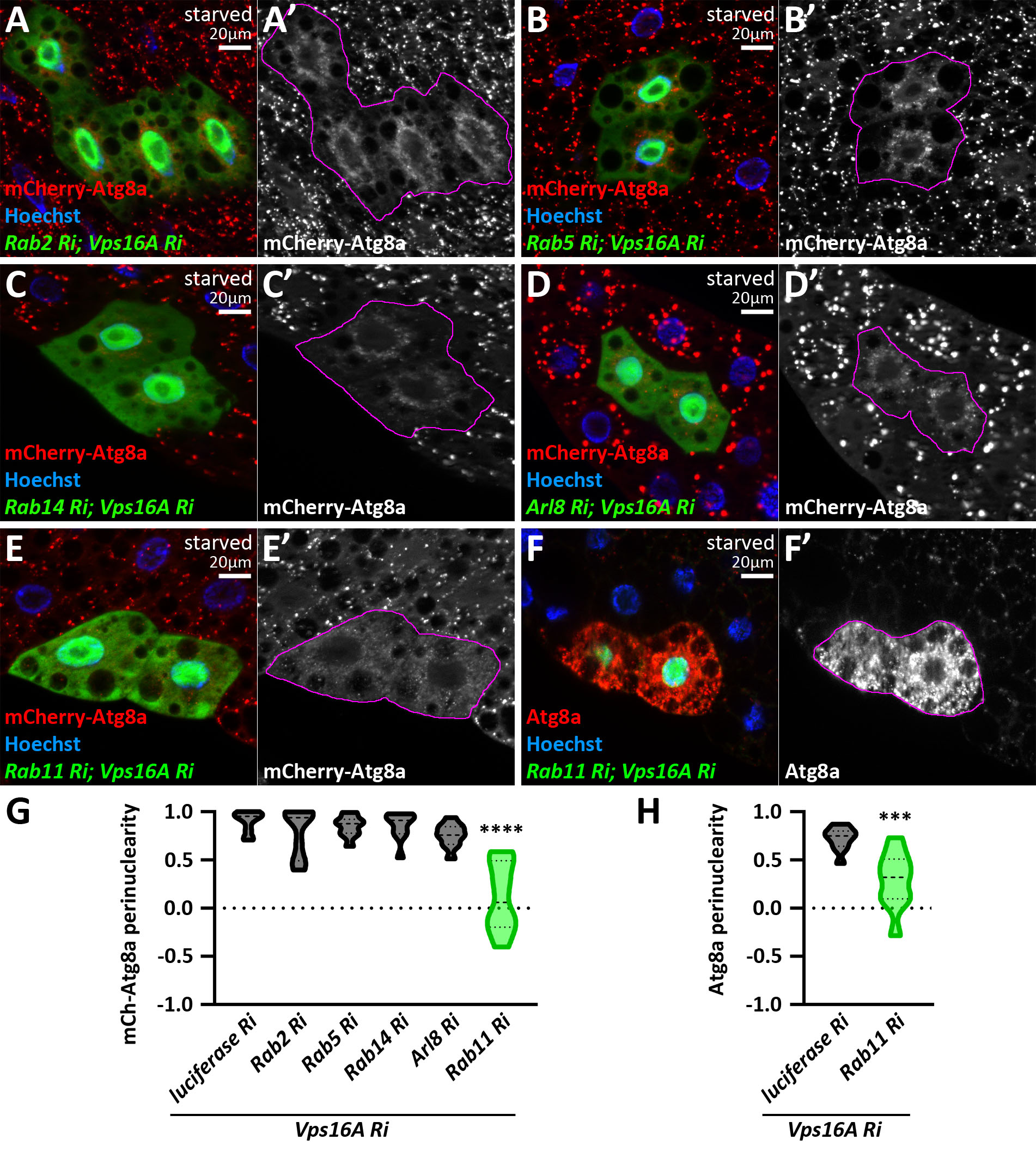

### Supplemental Data 5

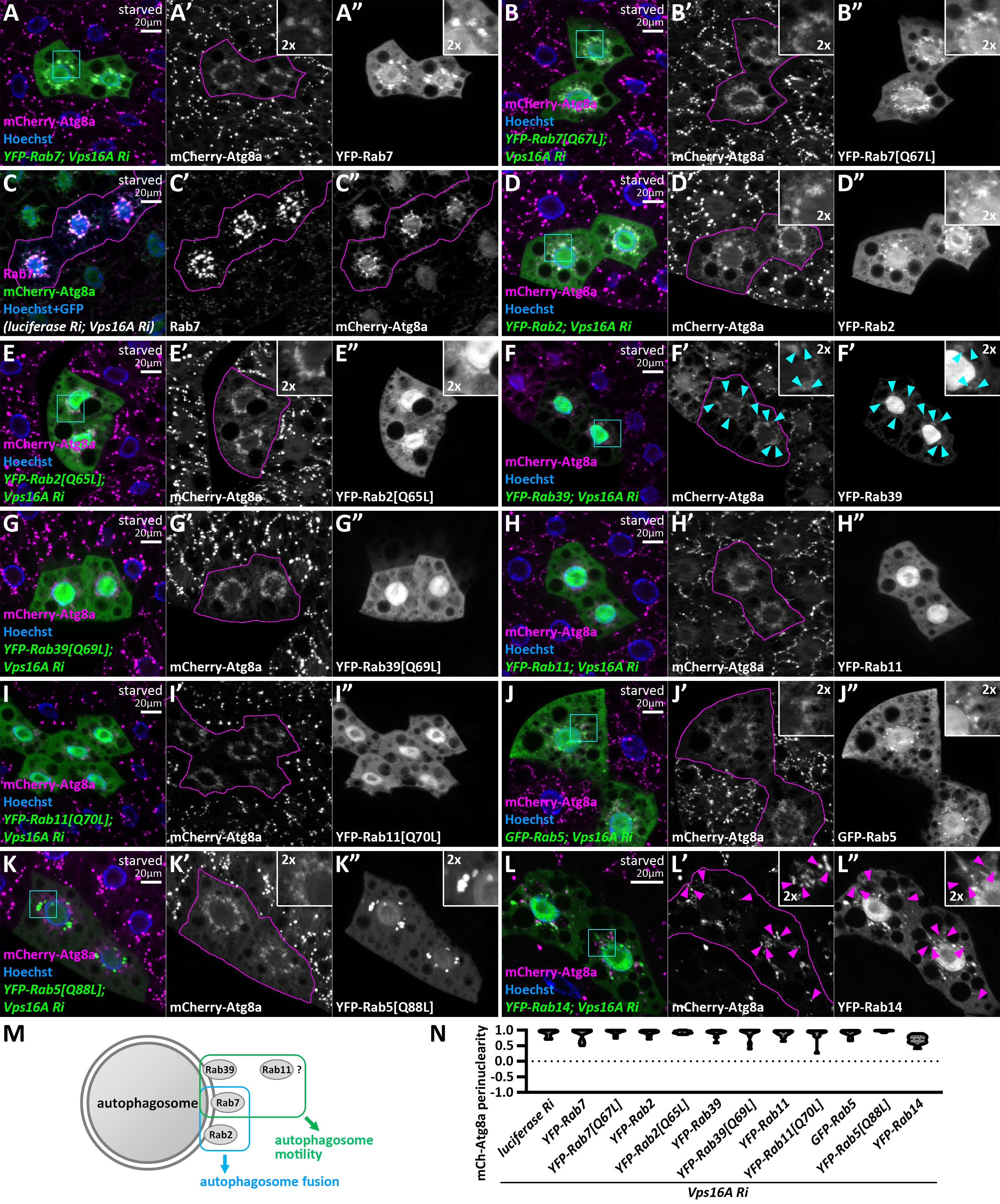

### Supplemental Data 6

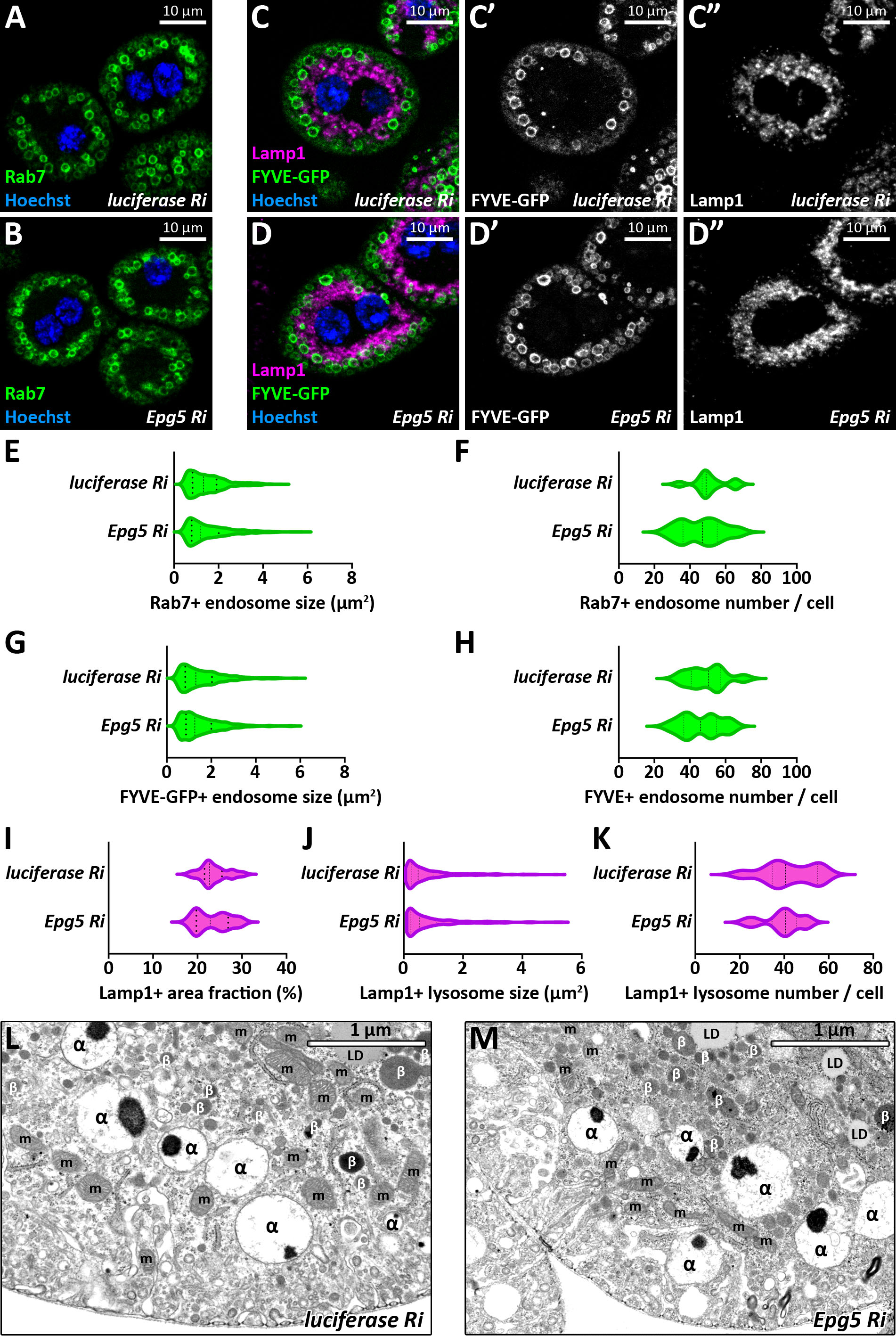

### Supplemental Data 7

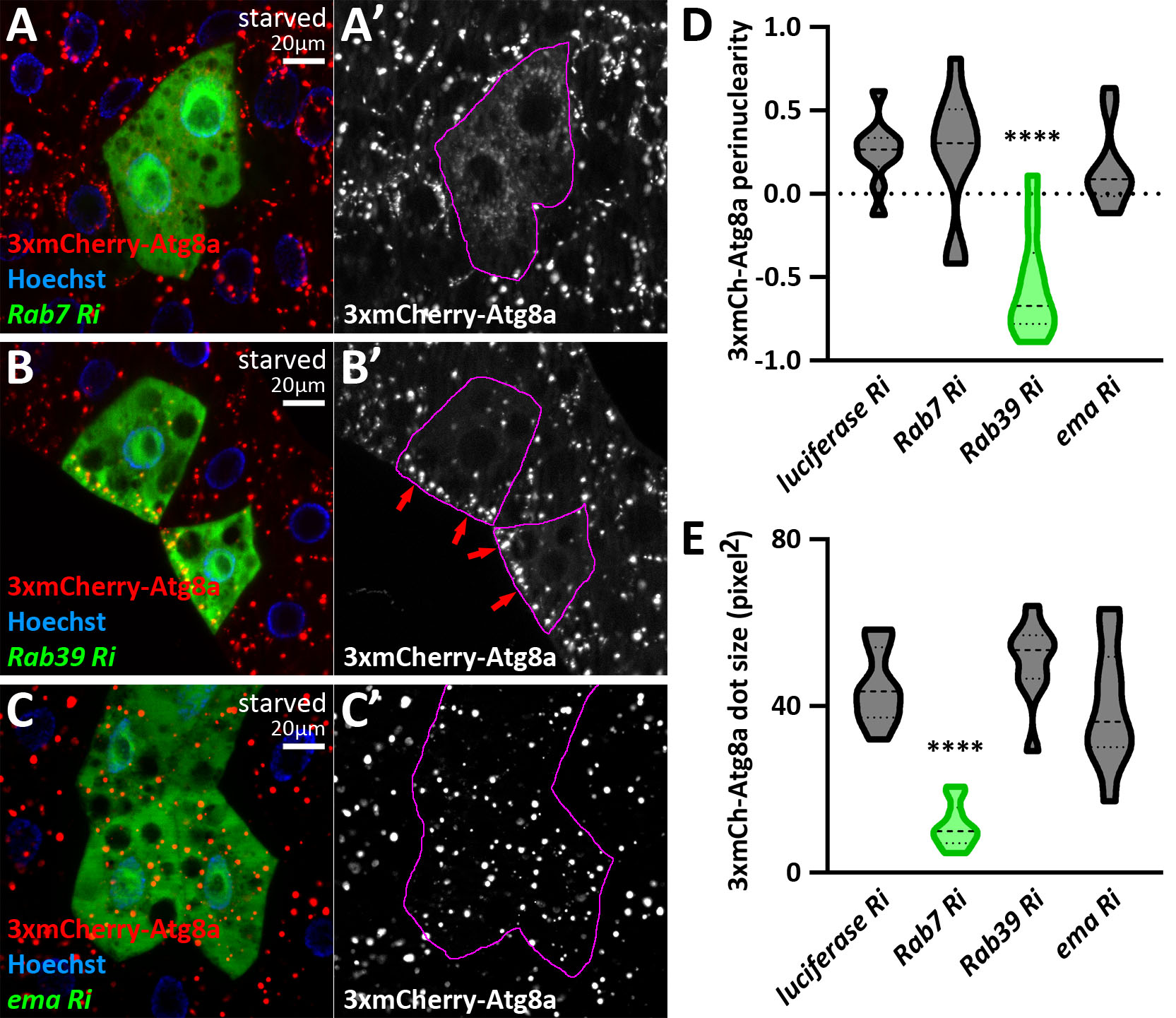

### Supplemental Data 8

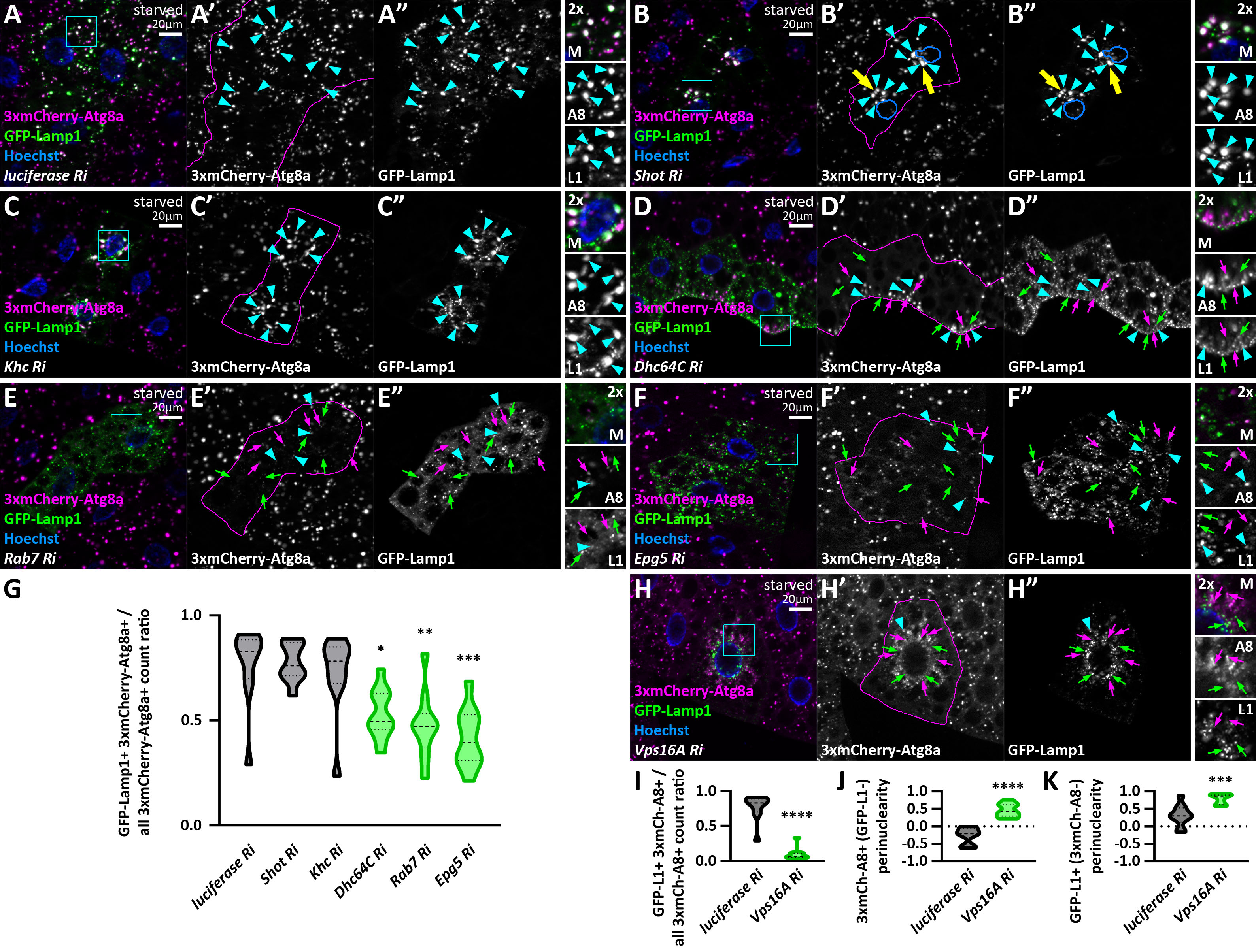

### Supplementary File 1

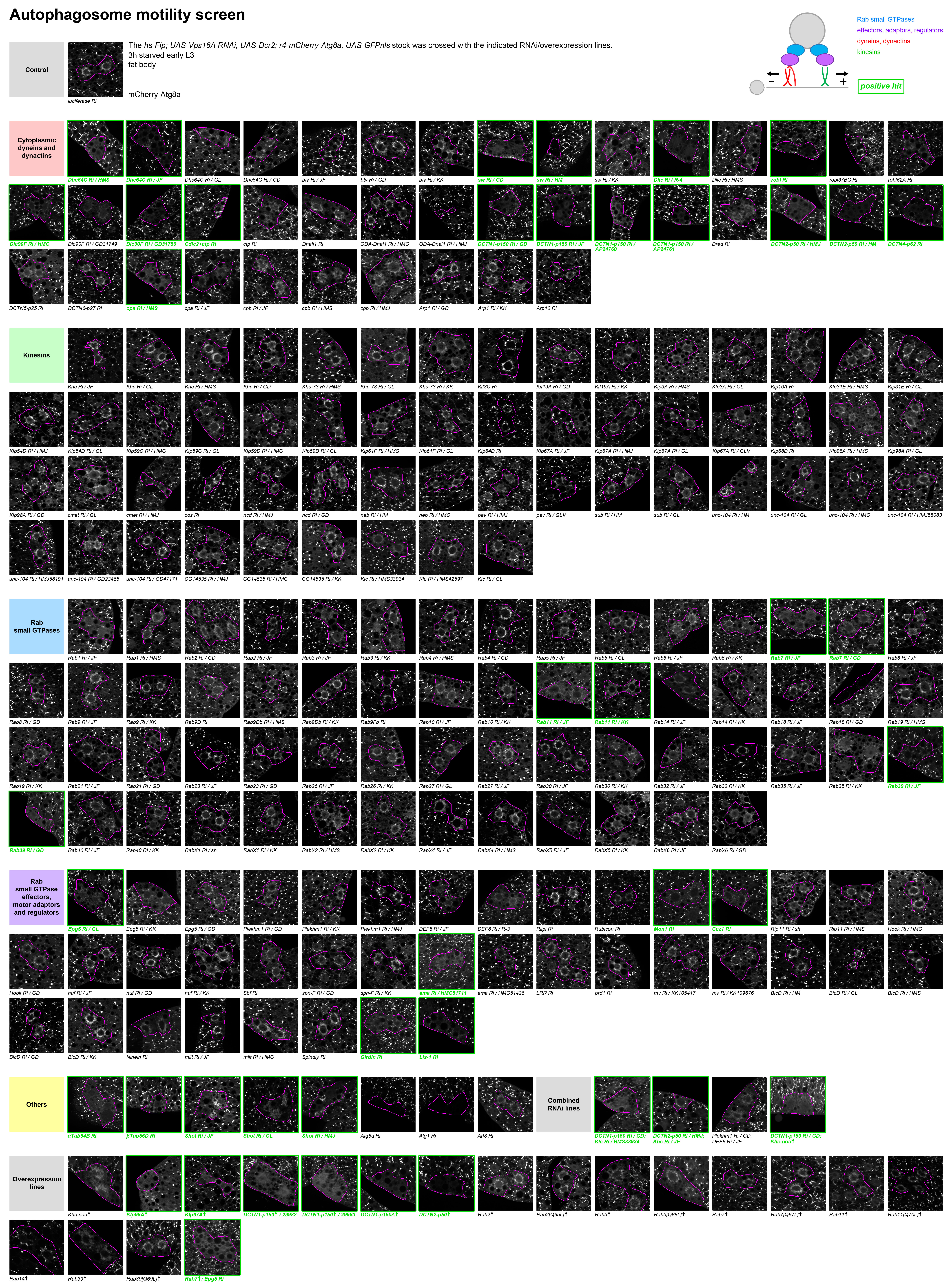
