## Supplementary Table 1 for "The Rab7-Epg5 and Rab39-ema modules cooperatively position autophagosomes for efficient lysosomal fusions"

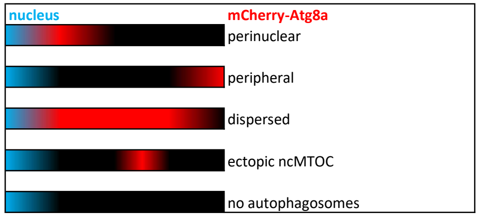


| RNAi | Stock # | Source | FlyBase ID | Type | Screen | Note |
|---|---|---|---|---|---|---|
| **control** | | | | | | |
| *luciferase RNAi JF* | 31603 | Bloomington Drosophila Stock Center | FBst0031603 | - | | |
| **Cytoplasmic dyneins and dynactins** | | | | | | |
| *Dhc64C RNAi HMS* | 36698 | Bloomington Drosophila Stock Center | FBst0036698 | DHC | | |
| *Dhc64C RNAi JF* | 28749 | Bloomington Drosophila Stock Center | FBst0028749 | DHC | | |
| *Dhc64C RNAi GL* | 36583 | Bloomington Drosophila Stock Center | FBst0036583 | DHC | | |
| *Dhc64C RNAi GD* | 28053 | Vienna Drosophila Resource Center | FBst0457266 | DHC | | |
| *btv RNAi JF* | 28373 | Bloomington Drosophila Stock Center | FBst0028373 | DHC | | |
| *btv RNAi GD* | 19323 | Vienna Drosophila Resource Center | FBst0453485 | DHC | | |
| *btv RNAi KK* | 105152 | Vienna Drosophila Resource Center | FBst0476980 | DHC | | |
| *sw RNAi GD* | 48334 | Vienna Drosophila Resource Center | FBst0467855 | DIC | | |
| *sw RNAi HM* | 30505 | Bloomington Drosophila Stock Center | FBst0030505 | DIC | | small cells |
| *sw RNAi KK* | 101559 | Vienna Drosophila Resource Center | FBst0473432 | DIC | | |
| *Dlic RNAi R-4* | 1938R-4 | Fly Stocks of National Institute of Genetics (Nig-Fly) | FBal0272979 | DLIC | | |
| *Dlic RNAi HMS* | 66981 | Bloomington Drosophila Stock Center | FBst0066981 | DLIC | | |
| *robl RNAi JF* | 31977 | Bloomington Drosophila Stock Center | FBst0031977 | DLC | | |
| *robl37BC RNAi HMC* | 64860 | Bloomington Drosophila Stock Center | FBst0064860 | DLC | | |
| *robl62A RNAi HMJ* | 54813 | Bloomington Drosophila Stock Center | FBst0054813 | DLC | | |
| *Dlc90F RNAi HMC* | 65189 | Bloomington Drosophila Stock Center | FBst0065189 | DLC | | |
| *Dlc90F RNAi GD* | 31749 | Vienna Drosophila Resource Center | FBst0459192 | DLC | | |
| *Dlc90F RNAi GD* | 31750 | Vienna Drosophila Resource Center | FBst0459194 | DLC | | |
| *Cdlc2+ctp RNAi HMS* | 42862 | Bloomington Drosophila Stock Center | FBst0042862 | DLC | | small cells |
| *ctp RNAi HMS* | 44044 | Bloomington Drosophila Stock Center | FBst0044044 | DLC | | |
| *Dnali1 RNAi HMS* | 63031 | Bloomington Drosophila Stock Center | FBst0063031 | DLC | | |
| *ODA-Dnal1 RNAi HMC* | 53295 | Bloomington Drosophila Stock Center | FBst0053295 | DLC | | |
| *ODA-Dnal1 RNAi HMJ* | 63590 | Bloomington Drosophila Stock Center | FBst0063590 | DLC | | small cells |
| *DCTN1-p150 RNAi GD* | 3785 | Vienna Drosophila Resource Center | FBst0462197 | dynactin | | |
| *DCTN1-p150 RNAi JF* | 27721 | Bloomington Drosophila Stock Center | FBst0027721 | dynactin | | |
| *DCTN1-p150 RNAi AP* | 24760 | Bloomington Drosophila Stock Center | FBst0024760 | dynactin | | |
| *DCTN1-p150 RNAi AP* | 24761 | Bloomington Drosophila Stock Center | FBst0024761 | dynactin | | |
| *Dred RNAi HMJ* | 50914 | Bloomington Drosophila Stock Center | FBst0050914 | dynactin | | |
| *DCTN2-p50 RNAi HMJ* | 63693 | Bloomington Drosophila Stock Center | FBst0063693 | dynactin | | |
| *DCTN2-p50 RNAi HM* | 28596 | Bloomington Drosophila Stock Center | FBst0028596 | dynactin | | |
| *DCTN4-p62 RNAi HMJ* | 60499 | Bloomington Drosophila Stock Center | FBst0060499 | dynactin | | |
| *DCTN5-p25 RNAi HM* | 30491 | Bloomington Drosophila Stock Center | FBst0030491 | dynactin | | |
| *DCTN6-p27 RNAi HMC* | 64679 | Bloomington Drosophila Stock Center | FBst0064679 | dynactin | | |
| *cpa RNAi HMS* | 41685 | Bloomington Drosophila Stock Center | FBst0041685 | dynactin | | |
| *cpa RNAi JF* | 31124 | Bloomington Drosophila Stock Center | FBst0031124 | dynactin | | |
| *cpb RNAi JF* | 26298 | Bloomington Drosophila Stock Center | FBst0026298 | dynactin | | |
| *cpb RNAi HMS* | 41952 | Bloomington Drosophila Stock Center | FBst0041952 | dynactin | | |
| *cpb RNAi HMJ* | 50954 | Bloomington Drosophila Stock Center | FBst0050954 | dynactin | | |
| *Arp1 RNAi GD* | 26007 | Vienna Drosophila Resource Center | FBst0456177 | dynactin | | |
| *Arp1 RNAi KK* | 100752 | Vienna Drosophila Resource Center | FBst0472625 | dynactin | | |
| *Arp10 RNAi HMC* | 64570 | Bloomington Drosophila Stock Center | FBst0064570 | dynactin | | |
| **Kinesins** | | | | | | |
| *Khc RNAi JF* | 25898 | Bloomington Drosophila Stock Center | FBst0025898 | kinesin motor | | small cells |
| *Khc RNAi GL* | 35409 | Bloomington Drosophila Stock Center | FBst0035409 | kinesin motor | | |
| *Khc RNAi HMS* | 35770 | Bloomington Drosophila Stock Center | FBst0035770 | kinesin motor | | |
| *Khc RNAi GD* | 44337 | Vienna Drosophila Resource Center | FBst0465523 | kinesin motor | | |
| *Khc-73 RNAi HMS* | 36733 | Bloomington Drosophila Stock Center | FBst0036733 | kinesin motor | | |

| | | | | | |
|---|---|---|---|---|---|
| *Khc-73 RNAi GL* | 38191 | Bloomington Drosophila Stock Center | FBst0038191 | kinesin motor | |
| *Khc-73 RNAi KK* | 105984 | Vienna Drosophila Resource Center | FBst0477810 | kinesin motor | |
| *Kif3C RNAi HMS* | 40886 | Bloomington Drosophila Stock Center | FBst0040886 | kinesin motor | |
| *Kif19A RNAi GD* | 36459 | Vienna Drosophila Resource Center | FBst0461698 | kinesin motor | |
| *Kif19A RNAi KK* | 106569 | Vienna Drosophila Resource Center | FBst0478393 | kinesin motor | |
| *Klp3A RNAi HMS* | 40944 | Bloomington Drosophila Stock Center | FBst0040944 | kinesin motor | |
| *Klp3A RNAi GL* | 43230 | Bloomington Drosophila Stock Center | FBst0043230 | kinesin motor | |
| *Klp10A RNAi GD* | 41534 | Vienna Drosophila Resource Center | FBst0464150 | kinesin motor | |
| *Klp31E RNAi HMS* | 40943 | Bloomington Drosophila Stock Center | FBst0040943 | kinesin motor | |
| *Klp31E RNAi GL* | 35473 | Bloomington Drosophila Stock Center | FBst0035473 | kinesin motor | |
| *Klp54D RNAi HMJ* | 63533 | Bloomington Drosophila Stock Center | FBst0063533 | kinesin motor | |
| *Klp54D RNAi GL* | 36577 | Bloomington Drosophila Stock Center | FBst0036577 | kinesin motor | |
| *Klp59C RNAi HMC* | 64673 | Bloomington Drosophila Stock Center | FBst0064673 | kinesin motor | |
| *Klp59C RNAi GL* | 35596 | Bloomington Drosophila Stock Center | FBst0035596 | kinesin motor | |
| *Klp59D RNAi HMC* | 64657 | Bloomington Drosophila Stock Center | FBst0064657 | kinesin motor | |
| *Klp59D RNAi GL* | 35474 | Bloomington Drosophila Stock Center | FBst0035474 | kinesin motor | |
| *Klp61F RNAi HMS* | 33685 | Bloomington Drosophila Stock Center | FBst0033685 | kinesin motor | |
| *Klp61F RNAi GL* | 35804 | Bloomington Drosophila Stock Center | FBst0035804 | kinesin motor | |
| *Klp64D RNAi HMS* | 40945 | Bloomington Drosophila Stock Center | FBst0040945 | kinesin motor | |
| *Klp67A RNAi JF* | 27549 | Bloomington Drosophila Stock Center | FBst0027549 | kinesin motor | small cells |
| *Klp67A RNAi HMJ* | 62383 | Bloomington Drosophila Stock Center | FBst0062383 | kinesin motor | |
| *Klp67A RNAi GL* | 35606 | Bloomington Drosophila Stock Center | FBst0035606 | kinesin motor | |
| *Klp67A RNAi GLV* | 36268 | Bloomington Drosophila Stock Center | FBst0036268 | kinesin motor | |
| *Klp68D RNAi JF* | 29410 | Bloomington Drosophila Stock Center | FBst0029410 | kinesin motor | |
| *Klp98A RNAi HMS* | 39037 | Bloomington Drosophila Stock Center | FBst0039037 | kinesin motor | |
| *Klp98A RNAi GL* | 50542 | Bloomington Drosophila Stock Center | FBst0050542 | kinesin motor | |
| *Klp98A RNAi GD* | 40603 | Vienna Drosophila Resource Center | FBst0463652 | kinesin motor | |
| *cmet RNAi GL* | 35816 | Bloomington Drosophila Stock Center | FBst0035816 | kinesin motor | |
| *cmet RNAi HMJ* | 54004 | Bloomington Drosophila Stock Center | FBst0054004 | kinesin motor | small cells |
| *cos RNAi HMC* | 44472 | Bloomington Drosophila Stock Center | FBst0044472 | kinesin motor | small cells |
| *ncd RNAi HMJ* | 58144 | Bloomington Drosophila Stock Center | FBst0058144 | kinesin motor | |
| *ncd RNAi GD* | 22570 | Vienna Drosophila Resource Center | FBst0454597 | kinesin motor | |
| *neb RNAi HM* | 28897 | Bloomington Drosophila Stock Center | FBst0028897 | kinesin motor | |
| *neb RNAi HMC* | 80389 | Bloomington Drosophila Stock Center | FBst0080389 | kinesin motor | |
| *pav RNAi HMJ* | 42573 | Bloomington Drosophila Stock Center | FBst0042573 | kinesin motor | |
| *pav RNAi GLV* | 35649 | Bloomington Drosophila Stock Center | FBst0035649 | kinesin motor | |
| *sub RNAi HM* | 28570 | Bloomington Drosophila Stock Center | FBst0028570 | kinesin motor | |
| *sub RNAi GL* | 36623 | Bloomington Drosophila Stock Center | FBst0036623 | kinesin motor | |
| *unc-104 RNAi HM* | 28951 | Bloomington Drosophila Stock Center | FBst0028951 | kinesin motor | small cells |
| *unc-104 RNAi GL* | 43264 | Bloomington Drosophila Stock Center | FBst0043264 | kinesin motor | |
| *unc-104 RNAi HMC* | 53296 | Bloomington Drosophila Stock Center | FBst0053296 | kinesin motor | |
| *unc-104 RNAi HMJ* | 58083 | Bloomington Drosophila Stock Center | FBst0058083 | kinesin motor | |
| *unc-104 RNAi HMJ* | 58191 | Bloomington Drosophila Stock Center | FBst0058191 | kinesin motor | |
| *unc-104 RNAi GD* | 23465 | Vienna Drosophila Resource Center | FBst0455027 | kinesin motor | |
| *unc-104 RNAi GD* | 47171 | Vienna Drosophila Resource Center | FBst0467113 | kinesin motor | |
| *CG14535 RNAi HMJ* | 62335 | Bloomington Drosophila Stock Center | FBst0062335 | kinesin motor | |
| *CG14535 RNAi HMC* | 51936 | Bloomington Drosophila Stock Center | FBst0051936 | kinesin motor | |
| *CG14535 RNAi KK* | 108308 | Vienna Drosophila Resource Center | FBst0480120 | kinesin motor | |
| *Klc RNAi HMS* | 33934 | Bloomington Drosophila Stock Center | FBst0033934 | KLC | |
| *Klc RNAi HMS* | 42597 | Bloomington Drosophila Stock Center | FBst0042597 | KLC | |
| *Klc RNAi GL* | 36795 | Bloomington Drosophila Stock Center | FBst0036795 | KLC | |
| **Rab small GTPases** | | | | | |
| *Rab1 RNAi JF* | 27299 | Bloomington Drosophila Stock Center | FBst0027299 | Rab small GTPase | |
| *Rab1 RNAi HMS* | 34670 | Bloomington Drosophila Stock Center | FBst0034670 | Rab small GTPase | |
| *Rab2 RNAi GD* | 34767 | Vienna Drosophila Resource Center | FBst0460794 | Rab small GTPase | |
| *Rab2 RNAi JF* | 28701 | Bloomington Drosophila Stock Center | FBst0028701 | Rab small GTPase | |
| *Rab3 RNAi JF* | 31691 | Bloomington Drosophila Stock Center | FBst0031691 | Rab small GTPase | |
| *Rab3 RNAi KK* | 100787 | Vienna Drosophila Resource Center | FBst0472660 | Rab small GTPase | |
| *Rab4 RNAi HMS* | 33757 | Bloomington Drosophila Stock Center | FBst0033757 | Rab small GTPase | |
| *Rab4 RNAi GD* | 24672 | Vienna Drosophila Resource Center | FBst0455536 | Rab small GTPase | |
| *Rab5 RNAi JF* | 30518 | Bloomington Drosophila Stock Center | FBst0030518 | Rab small GTPase | |
| *Rab5 RNAi GL* | 67877 | Bloomington Drosophila Stock Center | FBst0067877 | Rab small GTPase | |

| | | | | | |
|---|---|---|---|---|---|
| *Rab6 RNAi JF* | 27490 | Bloomington Drosophila Stock Center | FBst0027490 | Rab small GTPase | |
| *Rab6 RNAi KK* | 100774 | Vienna Drosophila Resource Center | FBst0472647 | Rab small GTPase | |
| *Rab7 RNAi JF* | 27051 | Bloomington Drosophila Stock Center | FBst0027051 | Rab small GTPase | |
| *Rab7 RNAi GD* | 40337 | Vienna Drosophila Resource Center | FBst0463506 | Rab small GTPase | |
| *Rab8 RNAi JF* | 27519 | Bloomington Drosophila Stock Center | FBst0027519 | Rab small GTPase | |
| *Rab8 RNAi GD* | 28092 | Vienna Drosophila Resource Center | FBst0457287 | Rab small GTPase | |
| *Rab9 RNAi JF* | 31688 | Bloomington Drosophila Stock Center | FBst0031688 | Rab small GTPase | |
| *Rab9 RNAi KK* | 107192 | Vienna Drosophila Resource Center | FBst0479014 | Rab small GTPase | |
| *Rab9D RNAi GD* | 49738 | Vienna Drosophila Resource Center | FBst0468633 | Rab small GTPase | |
| *Rab9Db RNAi HMS* | 38269 | Bloomington Drosophila Stock Center | FBst0038269 | Rab small GTPase | |
| *Rab9Db RNAi KK* | 109089 | Vienna Drosophila Resource Center | FBst0480875 | Rab small GTPase | |
| *Rab9Fb RNAi HMS* | 34374 | Bloomington Drosophila Stock Center | FBst0034374 | Rab small GTPase | small cells |
| *Rab10 RNAi JF* | 26289 | Bloomington Drosophila Stock Center | FBst0026289 | Rab small GTPase | |
| *Rab10 RNAi KK* | 101454 | Vienna Drosophila Resource Center | FBst0473327 | Rab small GTPase | |
| *Rab11 RNAi JF* | 27730 | Bloomington Drosophila Stock Center | FBst0027730 | Rab small GTPase | |
| *Rab11 RNAi KK* | 108382 | Vienna Drosophila Resource Center | FBst0480193 | Rab small GTPase | weaker than *Rab11 RNAi JF* |
| *Rab14 RNAi JF* | 28708 | Bloomington Drosophila Stock Center | FBst0028708 | Rab small GTPase | |
| *Rab14 RNAi KK* | 104392 | Vienna Drosophila Resource Center | FBst0476250 | Rab small GTPase | |
| *Rab18 RNAi JF* | 27665 | Bloomington Drosophila Stock Center | FBst0027665 | Rab small GTPase | |
| *Rab18 RNAi GD* | 40228 | Vienna Drosophila Resource Center | FBst0463448 | Rab small GTPase | less autophagosomes; small cells |
| *Rab19 RNAi HMS* | 34607 | Bloomington Drosophila Stock Center | FBst0034607 | Rab small GTPase | |
| *Rab19 RNAi KK* | 103653 | Vienna Drosophila Resource Center | FBst0475511 | Rab small GTPase | |
| *Rab21 RNAi JF* | 29403 | Bloomington Drosophila Stock Center | FBst0029403 | Rab small GTPase | |
| *Rab21 RNAi GD* | 32941 | Vienna Drosophila Resource Center | FBst0459849 | Rab small GTPase | |
| *Rab23 RNAi JF* | 28025 | Bloomington Drosophila Stock Center | FBst0028025 | Rab small GTPase | |
| *Rab23 RNAi GD* | 13147 | Vienna Drosophila Resource Center | FBst0450870 | Rab small GTPase | |
| *Rab26 RNAi JF* | 31177 | Bloomington Drosophila Stock Center | FBst0031177 | Rab small GTPase | |
| *Rab26 RNAi KK* | 101330 | Vienna Drosophila Resource Center | FBst0473203 | Rab small GTPase | |
| *Rab27 RNAi GL* | 50537 | Bloomington Drosophila Stock Center | FBst0050537 | Rab small GTPase | |
| *Rab27 RNAi JF* | 31887 | Bloomington Drosophila Stock Center | FBst0031887 | Rab small GTPase | |
| *Rab30 RNAi JF* | 31120 | Bloomington Drosophila Stock Center | FBst0031120 | Rab small GTPase | |
| *Rab30 RNAi KK* | 110230 | Vienna Drosophila Resource Center | FBst0481811 | Rab small GTPase | |
| *Rab32 RNAi JF* | 28002 | Bloomington Drosophila Stock Center | FBst0028002 | Rab small GTPase | |
| *Rab32 RNAi KK* | 104348 | Vienna Drosophila Resource Center | FBst0476206 | Rab small GTPase | |
| *Rab35 RNAi JF* | 28342 | Bloomington Drosophila Stock Center | FBst0028342 | Rab small GTPase | |
| *Rab35 RNAi KK* | 101361 | Vienna Drosophila Resource Center | FBst0473234 | Rab small GTPase | |
| *Rab39 RNAi JF* | 25953 | Bloomington Drosophila Stock Center | FBst0025953 | Rab small GTPase | |
| *Rab39 RNAi GD* | 31665 | Vienna Drosophila Resource Center | FBst0459139 | Rab small GTPase | small cells |
| *Rab40 RNAi JF* | 29579 | Bloomington Drosophila Stock Center | FBst0029579 | Rab small GTPase | |
| *Rab40 RNAi KK* | 110563 | Vienna Drosophila Resource Center | FBst0482129 | Rab small GTPase | |
| *RabX1 RNAi shRNA* | 330763 | Vienna Drosophila Resource Center | FBst0494112 | Rab small GTPase | |
| *RabX1 RNAi KK* | 103039 | Vienna Drosophila Resource Center | FBst0474901 | Rab small GTPase | |
| *RabX2 RNAi HMS* | 32360 | Bloomington Drosophila Stock Center | FBst0032360 | Rab small GTPase | |
| *RabX2 RNAi KK* | 103311 | Vienna Drosophila Resource Center | FBst0475170 | Rab small GTPase | |
| *RabX4 RNAi JF* | 28704 | Bloomington Drosophila Stock Center | FBst0028704 | Rab small GTPase | |
| *RabX4 RNAi HMS* | 44070 | Bloomington Drosophila Stock Center | FBst0044070 | Rab small GTPase | |
| *RabX5 RNAi JF* | 28045 | Bloomington Drosophila Stock Center | FBst0028045 | Rab small GTPase | |
| *RabX5 RNAi KK* | 103630 | Vienna Drosophila Resource Center | FBst0475488 | Rab small GTPase | |
| *RabX6 RNAi JF* | 26281 | Bloomington Drosophila Stock Center | FBst0026281 | Rab small GTPase | |
| *RabX6 RNAi GD* | 50817 | Vienna Drosophila Resource Center | FBst0469219 | Rab small GTPase | |
| **Small GTPase effectors, motor adaptors and regulators** | | | | | |
| *Epg5 RNAi GL* | 35624 | Bloomington Drosophila Stock Center | FBst0035624 | Rab7 interactor | |
| *Epg5 RNAi KK* | 110420 | Vienna Drosophila Resource Center | FBst0481992 | Rab7 interactor | sometimes dispersed |
| *Epg5 RNAi GD* | 17469 | Vienna Drosophila Resource Center | FBst0452724 | Rab7 interactor | |
| *Plekhm1 RNAi GD* | 42065 | Vienna Drosophila Resource Center | FBst0464428 | Rab7 interactor | |
| *Plekhm1 RNAi KK* | 108079 | Vienna Drosophila Resource Center | FBst0479891 | Rab7 interactor | |
| *Plekhm1 RNAi HMJ* | HMJ23550 | Fly Stocks of National Institute of Genetics (Nig-Fly) | FBst1084430 | Rab7 interactor | |
| *DEF8 RNAi JF* | 28312 | Bloomington Drosophila Stock Center | FBst0028312 | Rab7 interactor | sometimes weakly dispersed |
| *DEF8 RNAi R-3* | 11534R-3 | Fly Stocks of National Institute of Genetics (Nig-Fly) | FBal0270900 | Rab7 interactor | |
| *RilpL RNAi R-3* | 11448R-3 | Fly Stocks of National Institute of Genetics (Nig-Fly) | FBal0270872 | mammalian dynein activator; Rab7 interactor | |
| *Rubicon RNAi KK* | 109424 | Vienna Drosophila Resource Center | FBst0481113 | Rab7 interactor | |
| *Mon1 RNAi R-1* | 11926R-1 | Fly Stocks of National Institute of Genetics (Nig-Fly) | FBal0271032 | Rab7 GEF | |

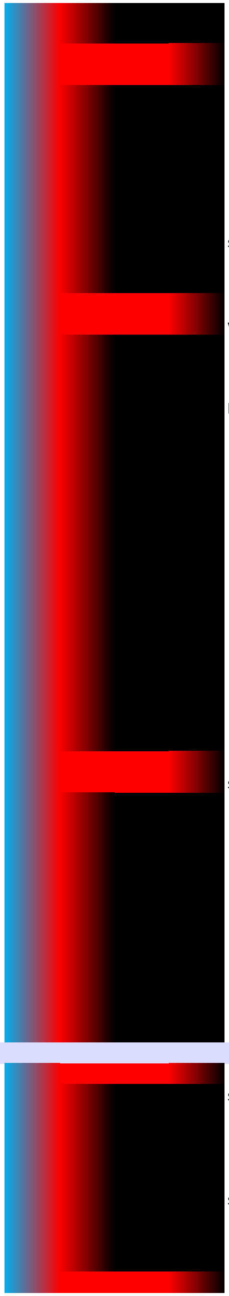

| | | | | | |
|---|---|---|---|---|---|
| *Ccz1 RNAi GD* | 18479 | Vienna Drosophila Resource Center | FBst0453144 | Rab7 GEF | |
| *Rip11 RNAi shRNA* | 330492 | Vienna Drosophila Resource Center | FBst0492207 | Rab11 interactor | |
| *Rip11 RNAi HMS* | 38325 | Bloomington Drosophila Stock Center | FBst0038325 | Rab11 interactor | |
| *Hook RNAi HMC* | 64663 | Bloomington Drosophila Stock Center | FBst0064663 | mammalian dynein activator; Rab11 interactor | |
| *Hook RNAi GD* | 35483 | Vienna Drosophila Resource Center | FBst0461178 | mammalian dynein activator; Rab11 interactor | |
| *nuf RNAi JF* | 31493 | Bloomington Drosophila Stock Center | FBst0031493 | mammalian dynein activator; Rab11 interactor | |
| *nuf RNAi GD* | 28069 | Vienna Drosophila Resource Center | FBst0457276 | mammalian dynein activator; Rab11 interactor | |
| *nuf RNAi KK* | 104172 | Vienna Drosophila Resource Center | FBst0476030 | mammalian dynein activator; Rab11 interactor | |
| *Sbf RNAi GD* | 22317 | Vienna Drosophila Resource Center | FBst0454518 | Rab11 interactor | |
| *spn-F RNAi GD* | 17015 | Vienna Drosophila Resource Center | FBst0452498 | Rab11 interactor | |
| *spn-F RNAi KK* | 107850 | Vienna Drosophila Resource Center | FBst0479663 | Rab11 interactor | |
| *ema RNAi HMC* | 51711 | Bloomington Drosophila Stock Center | FBst0051711 | Rab39 interactor | |
| *ema RNAi HMC* | 51426 | Bloomington Drosophila Stock Center | FBst0051426 | Rab39 interactor | |
| *LRR RNAi HMS* | 41686 | Bloomington Drosophila Stock Center | FBst0041686 | Rab39 interactor | |
| *prd1 RNAi KK* | 108557 | Vienna Drosophila Resource Center | FBst0480367 | Arl8 interactor | |
| *mv RNAi KK* | 105417 | Vienna Drosophila Resource Center | FBst0477244 | Rab2 interactor | |
| *mv RNAi KK* | 109676 | Vienna Drosophila Resource Center | FBst0481340 | Rab2 interactor | |
| *BicD RNAi HM* | 28571 | Bloomington Drosophila Stock Center | FBst0028571 | mammalian dynein activator; Rab2/Rab39 interactor | |
| *BicD RNAi GL* | 35405 | Bloomington Drosophila Stock Center | FBst0035405 | mammalian dynein activator; Rab2/Rab39 interactor | small cells |
| *BicD RNAi HMS* | 42929 | Bloomington Drosophila Stock Center | FBst0042929 | mammalian dynein activator; Rab2/Rab39 interactor | |
| *BicD RNAi GD* | 27683 | Vienna Drosophila Resource Center | FBst0457081 | mammalian dynein activator; Rab2/Rab39 interactor | |
| *BicD RNAi KK* | 108084 | Vienna Drosophila Resource Center | FBst0479896 | mammalian dynein activator; Rab2/Rab39 interactor | |
| *Ninein RNAi HMJ* | 62414 | Bloomington Drosophila Stock Center | FBst0062414 | mammalian dynein activator | |
| *milt RNAi JF* | 28385 | Bloomington Drosophila Stock Center | FBst0028385 | mammalian dynein activator | |
| *milt RNAi HMC* | 44477 | Bloomington Drosophila Stock Center | FBst0044477 | mammalian dynein activator | |
| *Spindly RNAi HMS* | 34933 | Bloomington Drosophila Stock Center | FBst0034933 | mammalian dynein activator | |
| *Girdin RNAi HMS* | 67960 | Bloomington Drosophila Stock Center | FBst0067960 | mammalian dynein activator candidate | |
| *Lis-1 RNAi JF* | 28663 | Bloomington Drosophila Stock Center | FBst0028663 | dynein regulator | |
| **Others** | | | | | |
| *αTub84B RNAi JF* | 31389 | Bloomington Drosophila Stock Center | FBst0031389 | microtubule subunit | |
| *βTub56D RNAi HMC* | 65028 | Bloomington Drosophila Stock Center | FBst0065028 | microtubule subunit | |
| *Shot RNAi JF* | 28336 | Bloomington Drosophila Stock Center | FBst0028336 | spectraplakin, MTOC component | |
| *Shot RNAi GL* | 41858 | Bloomington Drosophila Stock Center | FBst0041858 | spectraplakin, MTOC component | |
| *Shot RNAi HMJ* | 64041 | Bloomington Drosophila Stock Center | FBst0064041 | spectraplakin, MTOC component | |
| *Atg8a RNAi KK* | 109654 | Vienna Drosophila Resource Center | FBst0481318 | autophagosomal protein | |
| *Atg1 RNAi GL* | 35177 | Bloomington Drosophila Stock Center | FBst0035177 | autophagosome biogenesis regulator | |
| *Arl8 RNAi R-2* | 7891R-2 | Fly Stocks of National Institute of Genetics (Nig-Fly) | FBal0275763 | lysosomal small GTPase | |
| **Combined RNAi lines** | | | | | |
| *DCTN1-p150 RNAi GD; Klc RNAi HMS* | 785; 33934 | Vienna Drosophila Resource Center; Bloomington Drosophi | FBst0462197; | dynactin-kinesin double RNAi | |
| *DCTN2-p50 RNAi HMJ; Khc RNAi JF* | 693; 25898 | Bloomington Drosophila Stock Center | FBst0063693; | dynactin-kinesin double RNAi | sometimes a perinuclear and a peripheral subset |
| *Plekhm1 RNAi GD; DEF8 RNAi JF* | 065; 28312 | Vienna Drosophila Resource Center; Bloomington Drosophi | FBst0464428; | Rab7 interactor double RNAi | |
| *DCTN1-p150 RNAi GD; UAS-Khc-nod-LacZ* | 3785; 9912 | Vienna Drosophila Resource Center; Bloomington Drosophi | FBst0462197; | dynactin RNAi; minus end reporter | sometimes a peripheral subset |
| **Overexpression lines** | | | | | |
| *UAS-Khc-nod-LacZ* | 9912 | Bloomington Drosophila Stock Center | FBst0009912 | minus end reporter | not equally distributed around the nucleus |
| *UAS-Klp98A-3xHA* | F001727 | FlyORF | FBst0500838 | kinesin motor | |
| *UAS-Klp67A-3xHA* | F001232 | FlyORF | FBst0500800 | kinesin motor | |
| *UAS-GFP-DCTN1-p150* | 29982 | Bloomington Drosophila Stock Center | FBst0029982 | dynactin | sometimes peripheral |
| *UAS-GFP-DCTN1-p150* | 29983 | Bloomington Drosophila Stock Center | FBst0029983 | dynactin | sometimes peripheral |
| *UAS-DCTN1-p150Δ* | 51645 | Bloomington Drosophila Stock Center | FBst0051645 | dominant negative dynactin | |
| *UAS-DCTN2-p50* | 8784 | Bloomington Drosophila Stock Center | FBst0008784 | dynactin | |
| *UAS-YFP-Rab2* | 23246 | Bloomington Drosophila Stock Center | FBst0023246 | Rab small GTPase | on autophagosomes |
| *UAS-YFP-Rab2[Q65L]* | 9761 | Bloomington Drosophila Stock Center | FBst0009761 | constitutively active Rab small GTPase | on autophagosomes |
| *UAS-GFP-Rab5* | 43336 | Bloomington Drosophila Stock Center | FBst0043336 | Rab small GTPase | |
| *UAS-YFP-Rab5[Q88L]* | 9774 | Bloomington Drosophila Stock Center | FBst0009774 | constitutively active Rab small GTPase | |
| *UAS-YFP-Rab7* | 23641 | Bloomington Drosophila Stock Center | FBst0023641 | Rab small GTPase | on autophagosomes |
| *UAS-YFP-Rab7[Q67L]* | 24103 | Bloomington Drosophila Stock Center | FBst0024103 | constitutively active Rab small GTPase | on autophagosomes |
| *UAS-YFP-Rab11* | 50782 | Bloomington Drosophila Stock Center | FBst0050782 | Rab small GTPase | |
| *UAS-YFP-Rab11[Q70L]* | 23260 | Bloomington Drosophila Stock Center | FBst0023260 | constitutively active Rab small GTPase | |
| *UAS-YFP-Rab14* | 9793 | Bloomington Drosophila Stock Center | FBst0009793 | Rab small GTPase | |
| *UAS-YFP-Rab39* | 9825 | Bloomington Drosophila Stock Center | FBst0009825 | Rab small GTPase | on autophagosomes |
| *UAS-YFP-Rab39[Q69L]* | 9822 | Bloomington Drosophila Stock Center | FBst0009822 | constitutively active Rab small GTPase | |
| *UAS-YFP-Rab7; Epg5 RNAi GL* | 641; 35624 | Bloomington Drosophila Stock Center | FBst0023641; | Rab small GTPase; Rab7 interactor | on autophagosomes |
