## Supplementary Table 2 for "The Rab7-Epg5 and Rab39-ema modules cooperatively position autophagosomes for efficient lysosomal fusions"

| Figure | Panel | Genotype |
|---|---|---|
| **Figure 1** | B | *hs-Flp / +; UAS-DCR2, UAS-Vps16A RNAi / +; actin>CD2>Gal4, r4-mCherry-Atg8a, UAS-GFPnls / UAS-pLUC RNAi* |
| | C | *hs-Flp / +; UAS-DCR2, UAS-Vps16A RNAi / +; actin>CD2>Gal4, UAS-GFPnls / UAS-pLUC RNAi* |
| | D | *hs-Flp / +; UAS-DCR2, UAS-Vps16A RNAi / +; actin>CD2>Gal4, r4-mCherry-Atg8a, UAS-GFPnls / UAS-αTub84B RNAi* |
| | E | *hs-Flp / +; UAS-DCR2, UAS-Vps16A RNAi / UAS-βTub56D RNAi; actin>CD2>Gal4, r4-mCherry-Atg8a, UAS-GFPnls / +* |
| | J | *hs-Flp / +; UAS-DCR2, UAS-Vps16A RNAi / +; actin>CD2>Gal4, r4-mCherry-Atg8a, UAS-GFPnls / UAS-Shot RNAi/1* |
| | L | *hs-Flp / +; UAS-DCR2, UAS-Vps16A RNAi / +; actin>CD2>Gal4, UAS-GFPnls / UAS-Khc-nod-LacZ* |
| | M | *hs-Flp / hs-Flp; UAS-DCR2, UAS-Vps16A RNAi / UAS-DCR2, UAS-Vps16A RNAi; actin>CD2>Gal4, UAS-GFPnls / TM3* |
| | N | *hs-Flp / +; UAS-DCR2, UAS-Vps16A RNAi / +; actin>CD2>Gal4, UAS-GFPnls / UAS-Shot RNAi/1* |
| | O | *hs-Flp / +; UAS-DCR2 / UAS-Shot RNAi/3; actin>CD2>Gal4, UAS-GFPnls / UAS-Khc-nod-LacZ* |
| | P | *hs-Flp / +; UAS-DCR2, UAS-Vps16A RNAi / UAS-Shot RNAi/3; actin>CD2>Gal4, UAS-GFPnls / UAS-Khc-nod-LacZ* |
| **Figure 2** | A | *hs-Flp / +; UAS-DCR2, UAS-Vps16A RNAi / +; actin>CD2>Gal4, r4-mCherry-Atg8a, UAS-GFPnls / UAS-Dhc64C RNAi/1* |
| | B | *hs-Flp / +; UAS-DCR2, UAS-Vps16A RNAi / +; actin>CD2>Gal4, r4-mCherry-Atg8a, UAS-GFPnls / UAS-sw RNAi* |
| | C | *hs-Flp / +; UAS-DCR2, UAS-Vps16A RNAi / +; actin>CD2>Gal4, r4-mCherry-Atg8a, UAS-GFPnls / UAS-Dlic RNAi* |
| | D | *hs-Flp / +; UAS-DCR2, UAS-Vps16A RNAi / +; actin>CD2>Gal4, r4-mCherry-Atg8a, UAS-GFPnls / UAS-robl RNAi* |
| | E | *hs-Flp / +; UAS-DCR2, UAS-Vps16A RNAi / UAS-DCTN1-p150 RNAi/1; actin>CD2>Gal4, r4-mCherry-Atg8a, UAS-GFPnls / +* |
| | F | *hs-Flp / +; UAS-DCR2, UAS-Vps16A RNAi / UAS-DCTN2-p150 RNAi/1; actin>CD2>Gal4, r4-mCherry-Atg8a, UAS-GFPnls / +* |
| | G | *hs-Flp / +; UAS-DCR2, UAS-Vps16A RNAi / UAS-DCTN4-p62 RNAi; actin>CD2>Gal4, r4-mCherry-Atg8a, UAS-GFPnls / +* |
| | H | *hs-Flp / +; UAS-DCR2, UAS-Vps16A RNAi / +; actin>CD2>Gal4, r4-mCherry-Atg8a, UAS-GFPnls / UAS-Khc RNAi* |
| | I | *hs-Flp / +; UAS-DCR2, UAS-Vps16A RNAi / +; actin>CD2>Gal4, r4-mCherry-Atg8a, UAS-GFPnls / UAS-Klc RNAi* |
| **Figure 3** | A | *hs-Flp / +; UAS-DCR2, UAS-Vps16A RNAi / +; actin>CD2>Gal4, r4-mCherry-Atg8a, UAS-GFPnls / UAS-Klp98A-3xHA* |
| | B | *hs-Flp / +; UAS-DCR2, UAS-Vps16A RNAi / +; actin>CD2>Gal4, r4-mCherry-Atg8a, UAS-GFPnls / UAS-Klp67A-3xHA* |
| | C | *hs-Flp / +; UAS-DCR2, UAS-Vps16A RNAi / UAS-DCTN1-p150 RNAi/1; actin>CD2>Gal4, r4-mCherry-Atg8a, UAS-GFPnls / UAS-Klc RNAi* |
| | D | *hs-Flp / +; UAS-DCR2, UAS-Vps16A RNAi / +; actin>CD2>Gal4, r4-mCherry-Atg8a, UAS-GFPnls / UAS-Khc-nod-LacZ* |
| | E | *hs-Flp / +; UAS-DCR2, UAS-Vps16A RNAi / UAS-DCTN1-p150 RNAi/1; actin>CD2>Gal4, r4-mCherry-Atg8a, UAS-GFPnls / UAS-Khc-nod-LacZ* |
| | G | *hs-Flp / +; UAS-DCR2 / +; actin>CD2>Gal4, UAS-GFPnls / UAS-Klp98A-3xHA* |
| | H | *hs-Flp / +; UAS-DCR2, UAS-Vps16A RNAi / +; actin>CD2>Gal4, UAS-GFPnls / UAS-Klp98A-3xHA* |
| **Figure 4** | A | *hs-Flp / +; UAS-DCR2, UAS-Vps16A RNAi / +; actin>CD2>Gal4, r4-mCherry-Atg8a, UAS-GFPnls / UAS-Rab7 RNAi* |
| | B | *hs-Flp / +; UAS-DCR2, UAS-Vps16A RNAi / +; actin>CD2>Gal4, r4-mCherry-Atg8a, UAS-GFPnls / UAS-Epg5 RNAi* |
| | C | *hs-Flp / +; UAS-DCR2, UAS-Vps16A RNAi / +; actin>CD2>Gal4, r4-mCherry-Atg8a, UAS-GFPnls / UAS-Mon1 RNAi* |
| | D | *hs-Flp / +; UAS-DCR2, UAS-Vps16A RNAi / +; actin>CD2>Gal4, r4-mCherry-Atg8a, UAS-GFPnls / UAS-Ccz1 RNAi* |
| | E | *hs-Flp / +; UAS-DCR2, UAS-Vps16A RNAi / +; actin>CD2>Gal4, r4-mCherry-Atg8a, UAS-GFPnls / UAS-Rab39 RNAi/1* |
| | F | *hs-Flp / +; UAS-DCR2, UAS-Vps16A RNAi / UAS-ema RNAi; actin>CD2>Gal4, r4-mCherry-Atg8a, UAS-GFPnls / +* |
| | G | *hs-Flp / +; UAS-DCR2, UAS-Vps16A RNAi / UAS-Plekhm1 RNAi; actin>CD2>Gal4, r4-mCherry-Atg8a, UAS-GFPnls / +* |
| | H | *hs-Flp / +; UAS-DCR2, UAS-Vps16A RNAi / UAS-prd1 RNAi; actin>CD2>Gal4, r4-mCherry-Atg8a, UAS-GFPnls / +* |
| **Figure 5** | A | *hs-Flp / +; UAS-DCR2, UAS-Vps16A RNAi / +; actin>CD2>Gal4, UAS-GFPnls / UAS-Epg5 RNAi* |
| | B | *hs-Flp / +; UAS-DCR2, UAS-Vps16A RNAi / +; actin>CD2>Gal4, UAS-GFPnls / UAS-Epg5 RNAi* |
| | C | *hs-Flp / +; UAS-DCR2, UAS-Vps16A RNAi / UAS-YFP-Rab7; actin>CD2>Gal4, r4-mCherry-Atg8a, UAS-GFPnls / UAS-Epg5 RNAi* |
| | D | *hs-Flp / +; UAS-DCR2, UAS-Vps16A RNAi / +; actin>CD2>Gal4, UAS-GFPnls / UAS-Epg5 RNAi* |
| | I | *Epg5-9xHA* (S2R+ cells) |
| | J | *Epg5-9xHA* (S2R+ cells) |
| **Figure 6** | A | *hs-Flp / +; UAS-Snap29 RNAi / +; actin>CD2>Gal4, r4-mCherry-Atg8a, UAS-GFPnls / UAS-pLUC RNAi* |
| | B | *hs-Flp / +; UAS-Snap29 RNAi / +; actin>CD2>Gal4, r4-mCherry-Atg8a, UAS-GFPnls / UAS-Shot RNAi/2* |
| | C | *hs-Flp / +; UAS-Snap29 RNAi / +; actin>CD2>Gal4, r4-mCherry-Atg8a, UAS-GFPnls / UAS-Dhc64C RNAi/1* |
| | D | *hs-Flp / +; UAS-Snap29 RNAi / +; actin>CD2>Gal4, r4-mCherry-Atg8a, UAS-GFPnls / UAS-Khc RNAi* |
| | E | *hs-Flp / +; UAS-Snap29 RNAi / +; actin>CD2>Gal4, r4-mCherry-Atg8a, UAS-GFPnls / UAS-Rab7 RNAi* |
| | F | *hs-Flp / +; UAS-Snap29 RNAi / +; actin>CD2>Gal4, r4-mCherry-Atg8a, UAS-GFPnls / UAS-Epg5 RNAi* |
| | G | *hs-Flp / +; UAS-Snap29 RNAi / +; actin>CD2>Gal4, r4-mCherry-Atg8a, UAS-GFPnls / UAS-Rab39 RNAi/1* |
| | H | *hs-Flp / +; UAS-Snap29 RNAi / UAS-ema RNAi; actin>CD2>Gal4, r4-mCherry-Atg8a, UAS-GFPnls / +* |
| **Figure 7** | A | *hs-Flp / +; UAS-DCR2, UAS-Vps16A RNAi / +; actin>CD2>Gal4, UAS-GFPnls / UAS-pLUC RNAi* |
| | B | *hs-Flp / +; UAS-DCR2, UAS-Vps16A RNAi / +; actin>CD2>Gal4, UAS-GFPnls / UAS-Shot RNAi/1* |
| | C | *hs-Flp / +; UAS-DCR2, UAS-Vps16A RNAi / +; actin>CD2>Gal4, UAS-GFPnls / UAS-Epg5 RNAi* |
| | D | *hs-Flp / +; UAS-DCR2, UAS-Vps16A RNAi / +; actin>CD2>Gal4, UAS-GFPnls / UAS-Dhc64C RNAi/1* |
| | E | *hs-Flp / +; UAS-DCR2, UAS-Vps16A RNAi / +; actin>CD2>Gal4, UAS-GFPnls / UAS-Khc RNAi* |
| | F | *hs-Flp / +; UAS-DCR2, UAS-Vps16A RNAi / +; actin>CD2>Gal4, UAS-GFPnls / UAS-Rab7 RNAi* |
| | G | *hs-Flp / +; UAS-DCR2, UAS-Vps16A RNAi / +; actin>CD2>Gal4, UAS-GFPnls / UAS-Rab39 RNAi/1* |
| | H | *hs-Flp / +; UAS-DCR2, UAS-Vps16A RNAi / UAS-ema RNAi; actin>CD2>Gal4, UAS-GFPnls / +* |
| | I | *hs-Flp / +; UAS-DCR2, UAS-Vps16A RNAi / +; actin>CD2>Gal4, UAS-GFPnls / UAS-Rab2 RNAi* |
| | J | *hs-Flp / +; UAS-DCR2, UAS-Vps16A RNAi / +; actin>CD2>Gal4, UAS-GFPnls / UAS-Arl8 RNAi* |
| **Figure 8** | A | *hs-Flp / +; 3xmCherry-Atg8a, UAS-2xEGFP / +; actin>CD2>Gal4, UAS-DCR2 / UAS-pLUC RNAi* |
| | B | *hs-Flp / +; 3xmCherry-Atg8a, UAS-2xEGFP / +; actin>CD2>Gal4, UAS-DCR2 / UAS-Epg5 RNAi* |
| | C | *hs-Flp / +; UAS-DCR2 / +; actin>CD2>Gal4, UAS-GFPnls / UAS-Epg5 RNAi* |
| | D | *hs-Flp / +; UAS-DCR2 / +; actin>CD2>Gal4, UAS-GFPnls / UAS-Epg5 RNAi* |
| **Figure 9** | A | *hs-Flp / +; 3xmCherry-Atg8a, UAS-2xEGFP / +; actin>CD2>Gal4, UAS-DCR2 / UAS-Dhc64C RNAi/1* |
| | B | *hs-Flp / +; 3xmCherry-Atg8a, UAS-2xEGFP / +; actin>CD2>Gal4, UAS-DCR2 / UAS-sw RNAi* |
| | C | *hs-Flp / +; 3xmCherry-Atg8a, UAS-2xEGFP / +; actin>CD2>Gal4, UAS-DCR2 / UAS-Dlic RNAi* |
| | D | *hs-Flp / +; 3xmCherry-Atg8a, UAS-2xEGFP / +; actin>CD2>Gal4, UAS-DCR2 / UAS-robl RNAi* |
| | E | *hs-Flp / +; 3xmCherry-Atg8a, UAS-2xEGFP / UAS-DCTN1-p150 RNAi/1; actin>CD2>Gal4, UAS-DCR2 / +* |
| | F | *hs-Flp / +; 3xmCherry-Atg8a, UAS-2xEGFP / +; actin>CD2>Gal4, UAS-DCR2 / UAS-Khc RNAi* |
| | G | *hs-Flp / +; 3xmCherry-Atg8a, UAS-2xEGFP / +; actin>CD2>Gal4, UAS-DCR2 / UAS-Klc RNAi* |
| | H | *hs-Flp / +; 3xmCherry-Atg8a, UAS-2xEGFP / +; actin>CD2>Gal4, UAS-DCR2 / UAS-Shot RNAi/1* |
| **Figure 10** | A | *hs-Flp / +; 3xmCherry-Atg8a, UAS-2xEGFP / +; actin>CD2>Gal4, UAS-DCR2 / UAS-pLUC RNAi* |
| | B | *hs-Flp / +; 3xmCherry-Atg8a, UAS-2xEGFP / +; actin>CD2>Gal4, UAS-DCR2 / UAS-Shot RNAi/1* |
| | C | *hs-Flp / +; 3xmCherry-Atg8a, UAS-2xEGFP / +; actin>CD2>Gal4, UAS-DCR2 / UAS-Khc RNAi* |
| | D | *hs-Flp / +; 3xmCherry-Atg8a, UAS-2xEGFP / +; actin>CD2>Gal4, UAS-DCR2 / UAS-Dhc64C RNAi/1* |
| | E | *hs-Flp / +; 3xmCherry-Atg8a, UAS-2xEGFP / UAS-DCTN1-p150 RNAi/1; actin>CD2>Gal4, UAS-DCR2 / +* |
| | F | *hs-Flp / +; 3xmCherry-Atg8a, UAS-2xEGFP / +; actin>CD2>Gal4, UAS-DCR2 / UAS-Rab7 RNAi* |
| | G | *hs-Flp / +; 3xmCherry-Atg8a, UAS-2xEGFP / +; actin>CD2>Gal4, UAS-DCR2 / UAS-Epg5 RNAi* |
| **Figure 1—figure supplement 1** | A | *hs-Flp / +; UAS-DCR2, UAS-Vps16A RNAi / +; actin>CD2>Gal4, UAS-GFPnls / UAS-pLUC RNAi* |
| | B | *hs-Flp / +; UAS-DCR2, UAS-Vps16A RNAi / +; actin>CD2>Gal4, UAS-GFPnls / UAS-pLUC RNAi* |
| | C | *hs-Flp / hs-Flp; UAS-DCR2, UAS-Vps16A RNAi / UAS-DCR2, UAS-Vps16A RNAi; actin>CD2>Gal4, UAS-GFPnls / TM3* |
| | D | *hs-Flp / +; UAS-DCR2, UAS-Vps16A RNAi / UAS-Atg8a RNAi; actin>CD2>Gal4, r4-mCherry-Atg8a, UAS-GFPnls / +* |
| | E | *hs-Flp / +; UAS-DCR2, UAS-Vps16A RNAi / +; actin>CD2>Gal4, r4-mCherry-Atg8a, UAS-GFPnls / UAS-Atg1 RNAi* |
| | F | *hs-Flp / +; UAS-DCR2, UAS-Vps16A RNAi / +; actin>CD2>Gal4, r4-mCherry-Atg8a, UAS-GFPnls / UAS-Shot RNAi/2* |
| | G | *hs-Flp / +; UAS-DCR2, UAS-Vps16A RNAi / UAS-Shot RNAi/3; actin>CD2>Gal4, r4-mCherry-Atg8a, UAS-GFPnls / +* |
| **Figure 2—figure supplement 1** | A | *hs-Flp / +; UAS-DCR2, UAS-Vps16A RNAi / +; actin>CD2>Gal4, r4-mCherry-Atg8a, UAS-GFPnls / UAS-Dhc64C RNAi/2* |
| | B | *hs-Flp / +; UAS-DCR2, UAS-Vps16A RNAi / +; actin>CD2>Gal4, r4-mCherry-Atg8a, UAS-GFPnls / UAS-DCTN1-p150 RNAi/2* |
| | C | *hs-Flp / +; UAS-DCR2, UAS-Vps16A RNAi / +; actin>CD2>Gal4, r4-mCherry-Atg8a, UAS-GFPnls / UAS-DCTN2-p50 RNAi/2* |
| | D | *hs-Flp / +; UAS-DCR2, UAS-Vps16A RNAi / +; actin>CD2>Gal4, UAS-GFPnls / UAS-Dhc64C RNAi/1* |
| | E | *hs-Flp / +; UAS-DCR2, UAS-Vps16A RNAi / +; actin>CD2>Gal4, UAS-GFPnls / UAS-Dlic RNAi* |
| | F | *hs-Flp / +; UAS-DCR2, UAS-Vps16A RNAi / UAS-DCTN1-p150 RNAi/1; actin>CD2>Gal4, UAS-GFPnls / +* |
| | G | *hs-Flp / +; UAS-DCR2, UAS-Vps16A RNAi / +; actin>CD2>Gal4, UAS-GFPnls / UAS-Dhc64C RNAi/1* |
| | H | *hs-Flp / +; UAS-DCR2, UAS-Vps16A RNAi / +; actin>CD2>Gal4, UAS-GFPnls / UAS-Dlic RNAi* |

| | | |
|---|---|---|
| | I | *hs-Flp / +; UAS-DCR2, UAS-Vps16A RNAi / UAS-DCTN1-p150 RNAi/1; actin>CD2>Gal4, UAS-GFPnls / +* |
| | J | *hs-Flp / +; UAS-DCR2, UAS-Vps16A RNAi / UAS-DCTN1-p150 Δ ; actin>CD2>Gal4, r4-mCherry-Atg8a, UAS-GFPnls / +* |
| | K | *hs-Flp / +; UAS-DCR2, UAS-Vps16A RNAi / UAS-DCTN2-p50; actin>CD2>Gal4, r4-mCherry-Atg8a, UAS-GFPnls / +* |
| | L | *hs-Flp / +; UAS-DCR2, UAS-Vps16A RNAi / UAS-Dlc90F RNAi; actin>CD2>Gal4, r4-mCherry-Atg8a, UAS-GFPnls / +* |
| | M | *hs-Flp / +; UAS-DCR2, UAS-Vps16A RNAi / +; actin>CD2>Gal4, r4-mCherry-Atg8a, UAS-GFPnls / UAS-cpa RNAi* |
| | N | *hs-Flp / +; UAS-DCR2, UAS-Vps16A RNAi / UAS-Girdin RNAi; actin>CD2>Gal4, r4-mCherry-Atg8a, UAS-GFPnls / +* |
| | O | *hs-Flp / +; UAS-DCR2, UAS-Vps16A RNAi / +; actin>CD2>Gal4, r4-mCherry-Atg8a, UAS-GFPnls / UAS-Lis-1 RNAi* |
| | P | *hs-Flp / +; UAS-DCR2, UAS-Vps16A RNAi / +; actin>CD2>Gal4, UAS-GFPnls / UAS-Khc RNAi* |
| | Q | *hs-Flp / +; UAS-DCR2, UAS-Vps16A RNAi / +; actin>CD2>Gal4, UAS-GFPnls / UAS-Klc RNAi* |
| **Figure 4—figure supplement 1** | A | *hs-Flp / +; UAS-DCR2, UAS-Vps16A RNAi / UAS-Rab39 RNAi/2; actin>CD2>Gal4, r4-mCherry-Atg8a, UAS-GFPnls / +* |
| | B | *hs-Flp / +; UAS-DCR2, UAS-Vps16A RNAi / +; actin>CD2>Gal4, UAS-GFPnls / UAS-Rab7 RNAi* |
| | C | *hs-Flp / +; UAS-DCR2, UAS-Vps16A RNAi / +; actin>CD2>Gal4, UAS-GFPnls / UAS-Rab39 RNAi/1* |
| | D | *hs-Flp / +; UAS-DCR2, UAS-Vps16A RNAi / UAS-ema RNAi; actin>CD2>Gal4, UAS-GFPnls / +* |
| | E | *hs-Flp / +; UAS-DCR2, UAS-Vps16A RNAi / +; actin>CD2>Gal4, UAS-GFPnls / UAS-Rab7 RNAi* |
| | F | *hs-Flp / +; UAS-DCR2, UAS-Vps16A RNAi / +; actin>CD2>Gal4, UAS-GFPnls / UAS-Rab39 RNAi/1* |
| **Figure 4—figure supplement 2** | A | *hs-Flp / +; UAS-DCR2, UAS-Vps16A RNAi / +; actin>CD2>Gal4, r4-mCherry-Atg8a, UAS-GFPnls / UAS-Rab2 RNAi* |
| | B | *hs-Flp / +; UAS-DCR2, UAS-Vps16A RNAi / +; actin>CD2>Gal4, r4-mCherry-Atg8a, UAS-GFPnls / UAS-Rab5 RNAi* |
| | C | *hs-Flp / +; UAS-DCR2, UAS-Vps16A RNAi / +; actin>CD2>Gal4, r4-mCherry-Atg8a, UAS-GFPnls / UAS-Rab14 RNAi* |
| | D | *hs-Flp / +; UAS-DCR2, UAS-Vps16A RNAi / +; actin>CD2>Gal4, r4-mCherry-Atg8a, UAS-GFPnls / UAS-Arl8 RNAi* |
| | E | *hs-Flp / +; UAS-DCR2, UAS-Vps16A RNAi / +; actin>CD2>Gal4, r4-mCherry-Atg8a, UAS-GFPnls / UAS-Rab11 RNAi* |
| | F | *hs-Flp / +; UAS-DCR2, UAS-Vps16A RNAi / +; actin>CD2>Gal4, UAS-GFPnls / UAS-Rab11 RNAi* |
| **Figure 4—figure supplement 3** | A | *hs-Flp / +; UAS-DCR2, UAS-Vps16A RNAi / UAS-YFP-Rab7; actin>CD2>Gal4, r4-mCherry-Atg8a, UAS-GFPnls / +* |
| | B | *hs-Flp / +; UAS-DCR2, UAS-Vps16A RNAi / UAS-YFP-Rab7[Q67L]; actin>CD2>Gal4, r4-mCherry-Atg8a, UAS-GFPnls / +* |
| | C | *hs-Flp / +; UAS-DCR2, UAS-Vps16A RNAi / +; actin>CD2>Gal4, r4-mCherry-Atg8a, UAS-GFPnls / UAS-pLUC RNAi* |
| | D | *hs-Flp / +; UAS-DCR2, UAS-Vps16A RNAi / +; actin>CD2>Gal4, r4-mCherry-Atg8a, UAS-GFPnls / UAS-YFP-Rab2* |
| | E | *hs-Flp / +; UAS-DCR2, UAS-Vps16A RNAi / UAS-YFP-Rab2[Q65L]; actin>CD2>Gal4, r4-mCherry-Atg8a, UAS-GFPnls / +* |
| | F | *hs-Flp / +; UAS-DCR2, UAS-Vps16A RNAi / UAS-YFP-Rab39; actin>CD2>Gal4, r4-mCherry-Atg8a, UAS-GFPnls / +* |
| | G | *hs-Flp / UAS-YFP-Rab39[Q69L]; UAS-DCR2, UAS-Vps16A RNAi / +; actin>CD2>Gal4, r4-mCherry-Atg8a, UAS-GFPnls / +* |
| | H | *hs-Flp / +; UAS-DCR2, UAS-Vps16A RNAi / UAS-YFP-Rab11; actin>CD2>Gal4, r4-mCherry-Atg8a, UAS-GFPnls / +* |
| | I | *hs-Flp / +; UAS-DCR2, UAS-Vps16A RNAi / UAS-YFP-Rab11[Q70L]; actin>CD2>Gal4, r4-mCherry-Atg8a, UAS-GFPnls / +* |
| | J | *hs-Flp / +; UAS-DCR2, UAS-Vps16A RNAi / +; actin>CD2>Gal4, r4-mCherry-Atg8a, UAS-GFPnls / UAS-GFP-Rab5* |
| | K | *hs-Flp / +; UAS-DCR2, UAS-Vps16A RNAi / UAS-YFP-Rab5[Q88L]; actin>CD2>Gal4, r4-mCherry-Atg8a, UAS-GFPnls / +* |
| | L | *hs-Flp / +; UAS-DCR2, UAS-Vps16A RNAi / UAS-YFP-Rab14; actin>CD2>Gal4, r4-mCherry-Atg8a, UAS-GFPnls / +* |
| **Figure 5—figure supplement 1** | A | *UAS-DCR2 / +; prospero-Gal4 / UAS-pLUC RNAi* |
| | B | *UAS-DCR2 / +; prospero-Gal4 / UAS-Epg5 RNAi* |
| | C | *UAS-GFP-myc-2xFYVE / +; prospero-Gal4 / UAS-pLUC RNAi* |
| | D | *UAS-GFP-myc-2xFYVE / +; prospero-Gal4 / UAS-Epg5 RNAi* |
| | L | *UAS-DCR2 / +; prospero-Gal4 / UAS-pLUC RNAi* |
| | M | *UAS-DCR2 / +; prospero-Gal4 / UAS-Epg5 RNAi* |
| **Figure 9—figure supplement 1** | A | *hs-Flp / +; 3xmCherry-Atg8a, UAS-2xEGFP / +; actin>CD2>Gal4, UAS-DCR2 / UAS-Rab7 RNAi* |
| | B | *hs-Flp / +; 3xmCherry-Atg8a, UAS-2xEGFP / +; actin>CD2>Gal4, UAS-DCR2 / UAS-Rab39 RNAi/1* |
| | C | *hs-Flp / +; 3xmCherry-Atg8a, UAS-2xEGFP / UAS-ema RNAi; actin>CD2>Gal4, UAS-DCR2 / +* |
| **Figure 10—figure supplement 1** | A | *hs-Flp / +; 3xmCherry-Atg8a, UAS-GFP-Lamp1 / +; actin>CD2>Gal4, UAS-DCR2 / UAS-pLUC RNAi* |
| | B | *hs-Flp / +; 3xmCherry-Atg8a, UAS-GFP-Lamp1 / +; actin>CD2>Gal4, UAS-DCR2 / UAS-Shot RNAi/1* |
| | C | *hs-Flp / +; 3xmCherry-Atg8a, UAS-GFP-Lamp1 / +; actin>CD2>Gal4, UAS-DCR2 / UAS-Khc RNAi* |
| | D | *hs-Flp / +; 3xmCherry-Atg8a, UAS-GFP-Lamp1 / +; actin>CD2>Gal4, UAS-DCR2 / UAS-Dhc64C RNAi/1* |
| | E | *hs-Flp / +; 3xmCherry-Atg8a, UAS-GFP-Lamp1 / +; actin>CD2>Gal4, UAS-DCR2 / UAS-Rab7 RNAi* |
| | F | *hs-Flp / +; 3xmCherry-Atg8a, UAS-GFP-Lamp1 / +; actin>CD2>Gal4, UAS-DCR2 / UAS-Epg5 RNAi* |
| | H | *hs-Flp / +; 3xmCherry-Atg8a, UAS-GFP-Lamp1 / UAS-Vps16A RNAi; actin>CD2>Gal4, UAS-DCR2 / +* |

**Stocks used in the Figures**

| Stock | Stock # | Source | FlyBase ID |
|---|---|---|---|
| **Figures** | | | |
| *UAS-pLUC RNAi JF* | 31603 | Bloomington Drosophila Stock Center | FBst0031603 |
| *UAS-Vps16A RNAi GD* | 23769 | Vienna Drosophila Resource Center | FBst0455191 |
| *UAS-αTub84B RNAi JF* | 31389 | Bloomington Drosophila Stock Center | FBst0031389 |
| *UAS-βTub56D RNAi HMC* | 65028 | Bloomington Drosophila Stock Center | FBst0065028 |
| *UAS-Shot RNAi/1 JF* | 28336 | Bloomington Drosophila Stock Center | FBst0028336 |
| *UAS-Shot RNAi/2 GL* | 41858 | Bloomington Drosophila Stock Center | FBst0041858 |
| *UAS-Shot RNAi/3 HMJ* | 64041 | Bloomington Drosophila Stock Center | FBst0064041 |
| *UAS-Khc-nod-LacZ* | 9912 | Bloomington Drosophila Stock Center | FBst0009912 |
| *UAS-Dhc64C RNAi/1 HMS* | 36698 | Bloomington Drosophila Stock Center | FBst0036698 |
| *UAS-Dhc64C RNAi/2 JF* | 28749 | Bloomington Drosophila Stock Center | FBst0028749 |
| *UAS-sw RNAi GD* | 48334 | Vienna Drosophila Resource Center | FBst0467855 |
| *UAS-Dlic RNAi R-4* | 1938R-4 | Fly Stocks of National Institute of Genetics (Nig-Fly) | FBal0272979 |
| *UAS-robl RNAi JF* | 31977 | Bloomington Drosophila Stock Center | FBst0031977 |
| *UAS-DCTN1-p150 RNAi/1 GD* | 3785 | Vienna Drosophila Resource Center | FBst0462197 |
| *UAS-DCTN1-p150 RNAi/2 JF* | 27721 | Bloomington Drosophila Stock Center | FBst0027721 |
| *UAS-DCTN2-p50 RNAi/1 HMJ* | 63693 | Bloomington Drosophila Stock Center | FBst0063693 |
| *UAS-DCTN2-p50 RNAi/2 HM* | 28596 | Bloomington Drosophila Stock Center | FBst0028596 |
| *UAS-DCTN4-p62 RNAi HMJ* | 60499 | Bloomington Drosophila Stock Center | FBst0060499 |
| *UAS-Khc RNAi JF* | 25898 | Bloomington Drosophila Stock Center | FBst0025898 |
| *UAS-Klc RNAi HMS* | 33934 | Bloomington Drosophila Stock Center | FBst0033934 |
| *UAS-Klp98A-3xHA* | F001727 | FlyORF | FBst0500838 |
| *UAS-Klp67A-3xHA* | F001232 | FlyORF | FBst0500800 |
| *UAS-Rab7 RNAi JF* | 27051 | Bloomington Drosophila Stock Center | FBst0027051 |
| *UAS-Epg5 RNAi GL* | 35624 | Bloomington Drosophila Stock Center | FBst0035624 |
| *UAS-Mon1 RNAi R-1* | 11926R-1 | Fly Stocks of National Institute of Genetics (Nig-Fly) | FBal0271032 |
| *UAS-Ccz1 RNAi GD* | 18479 | Vienna Drosophila Resource Center | FBst0453144 |
| *UAS-Rab39 RNAi/1 JF* | 25953 | Bloomington Drosophila Stock Center | FBst0025953 |
| *UAS-Rab39 RNAi/2 GD* | 31665 | Vienna Drosophila Resource Center | FBst0459139 |
| *UAS-ema RNAi HMC* | 51711 | Bloomington Drosophila Stock Center | FBst0051711 |
| *UAS-Plekhm1 RNAi GD* | 42065 | Vienna Drosophila Resource Center | FBst0464428 |
| *UAS-prd1 RNAi KK* | 108557 | Vienna Drosophila Resource Center | FBst0480367 |
| *UAS-Atg8a RNAi KK* | 109654 | Vienna Drosophila Resource Center | FBst0481318 |
| *UAS-Atg1 RNAi GL* | 35177 | Bloomington Drosophila Stock Center | FBst0035177 |
| *UAS-DCTN1-p150Δ* | 51645 | Bloomington Drosophila Stock Center | FBst0051645 |
| *UAS-DCTN2-p50* | 8784 | Bloomington Drosophila Stock Center | FBst0008784 |
| *UAS-Dlc90F RNAi HMC* | 65189 | Bloomington Drosophila Stock Center | FBst0065189 |
| *UAS-cpa RNAi HMS* | 41685 | Bloomington Drosophila Stock Center | FBst0041685 |
| *UAS-Girdin RNAi HMS* | 67960 | Bloomington Drosophila Stock Center | FBst0067960 |
| *UAS-Lis-1 RNAi JF* | 28663 | Bloomington Drosophila Stock Center | FBst0028663 |
| *UAS-Rab2 RNAi GD* | 34767 | Vienna Drosophila Resource Center | FBst0460794 |

| | | | |
|---|---|---|---|
| *UAS-Rab5 RNAi JF* | 30518 | Bloomington Drosophila Stock Center | FBst0030518 |
| *UAS-Rab14 RNAi JF* | 28708 | Bloomington Drosophila Stock Center | FBst0028708 |
| *UAS-Arl8 RNAi R-2* | 7891R-2 | Fly Stocks of National Institute of Genetics (Nig-Fly) | FBal0275763 |
| *UAS-Rab11 RNAi JF* | 27730 | Bloomington Drosophila Stock Center | FBst0027730 |
| *UAS-YFP-Rab7* | 23641 | Bloomington Drosophila Stock Center | FBst0023641 |
| *UAS-YFP-Rab7[Q67L]* | 24103 | Bloomington Drosophila Stock Center | FBst0024103 |
| *UAS-YFP-Rab2* | 23246 | Bloomington Drosophila Stock Center | FBst0023246 |
| *UAS-YFP-Rab2[Q65L]* | 9761 | Bloomington Drosophila Stock Center | FBst0009761 |
| *UAS-YFP-Rab39* | 9825 | Bloomington Drosophila Stock Center | FBst0009825 |
| *UAS-YFP-Rab39[Q69L]* | 9822 | Bloomington Drosophila Stock Center | FBst0009822 |
| *UAS-YFP-Rab11* | 50782 | Bloomington Drosophila Stock Center | FBst0050782 |
| *UAS-YFP-Rab11[Q70L]* | 23260 | Bloomington Drosophila Stock Center | FBst0023260 |
| *UAS-GFP-Rab5* | 43336 | Bloomington Drosophila Stock Center | FBst0043336 |
| *UAS-YFP-Rab5[Q88L]* | 9774 | Bloomington Drosophila Stock Center | FBst0009774 |
| *UAS-YFP-Rab14* | 9793 | Bloomington Drosophila Stock Center | FBst0009793 |
| *UAS-Snap29 RNAi HMC* | 51893 | Bloomington Drosophila Stock Center | FBst0051893 |
